# ADAR1 regulates alternative splicing through an RNA editing-independent mechanism

**DOI:** 10.1101/2025.03.12.642916

**Authors:** Eduardo A. Sagredo, Victor Karlström, Alejandro Blanco, Paloma Moraga, Matias Vergara, Aino I. Jarvelin, Neus Visa, Katherine Marcelain, Alfredo Castello, Ricardo Armisén

## Abstract

Adenosine-to-inosine (A-to-I) RNA editing and alternative splicing occur co-transcriptionally and can regulate and influence each other. The RNA editing enzymes ADAR1 and ADAR2 catalyze A-to-I RNA editing, where it has been shown that expression changes in both RNA editing enzymes exert alternative splicing. ADAR1 manipulation has a significant impact on alternative splicing. While many of those changes are related to changes in their A-to-I RNA editing activity, we speculate that ADAR1 may also influence splicing in an editing-independent manner. In this work, the protein-protein interactome of ADAR1 revealed that ADAR1 interacts with spliceosome co-factors and auxiliary splicing regulators. We confirmed that ADAR1 does not only influence splicing through editing but also throughout editing-independent functions with even greater penetrance. We show that ADAR1 modulates the splicing of transcripts encoding splicing factors, with many of these splice sites overlapping with known splicing changes induced by the splicing factor ACIN1, suggesting indirect effects upon ADAR1 expression. In summary, we show that ADAR1 can regulate splicing in an editing-independent manner, which likely occurs by widespread alteration of the splicing factor isoform landscape.

## Introduction

The adenosine deaminase acting on RNA (ADAR) enzymes catalyze adenosine-to-inosine (A-to-I) RNA editing, one of the most prevalent RNA modifications in mammals ^1–3^. Inosine is recognized as guanosine by the cellular machinery, which can cause profound effects on RNA structure and translational fidelity^4–6^. The ADAR protein family consists of three members (ADAR1-3), two of which are catalytically active (ADAR1 and ADAR2)^3^. All ADAR enzymes contain two to three double-stranded RNA binding domains and a deamination domain^7^. ADAR1 is ubiquitously expressed and has two isoforms, ADAR1p110, which is primarily nuclear and nucleolar, while ADAR1p150 is induced by interferon and can shuttle between the nucleus and the cytoplasm^8,9^. Changes in ADAR1 expression and activity are associated with different diseases, including amyotrophic lateral sclerosis^10^, systemic lupus erythematosus^11^, and several types of cancers^12^. Recent data suggest a broad role of ADAR proteins in cell fate, showing that they can modify mRNA splicing sites^13^ and alternative polyadenylation sites^14^ and affect miRNA processing/maturation^15,16^ and RNA stability^17–19^ by altering canonical RNA-RNA and RNA-proteins interactions^20^. Several studies have established how RNA editing may influence alternative splicing by modifying splicing regulatory elements or by altering dsRNA structures. These alterations may affect the affinity of the splicing machinery and trans-acting factors for those regions, leading to alterations in splicing efficiency^13,21–24^. One of the most well-characterized RNA editing-dependent splicing events involves the glutamate receptor subunit 2 (GRIA2) transcript^25^, where a single recoding site of the GRIA2 pre-mRNA, in the amino acid 607 (known as Q/R site), is required for the excision of the intron 11, regulating the processing and proper mRNA cytoplasmic localization^26–28^. Moreover, ADAR2 can edit its transcript, generating a new splice site resulting in a transcript variant that encodes a truncated non-functional protein^27,29^. As virtually all mRNAs are spliced and a vast majority undergo RNA editing co-transcriptionally, it has been proposed that these two processes can mutually influence each other. Indeed, transcriptomic analyses revealed that ADAR1 and ADAR2 expression alterations significantly affect exon usage and alternative splicing in animal and cellular models ^13,21,30,31^. In counterpart, it has been shown that the pharmacological inhibition of the spliceosome influences ADAR editing function in those sites that compromise an intron or other RNA structures that can enhance RNA editing due to secondary structures^21,31^.

ADAR1 knock-out mice die at early embryonic stages due to widespread apoptosis^32–34^. This phenotype can be rescued to a varying extent by adding back catalytically inactive ADAR1 mutants, indicating that A-to-I editing is not the only role of ADAR1 in RNA metabolism^35,36^. In agreement, it has been shown that ADAR can interact with other RBPs and influence their activity, further reinforcing the idea of an editing-independent function^20^. In the early days of genomic studies, Solomon et al. (2013)^13^ intersected alternative splicing and RNA editing analyses upon ADAR1 knockdown, showing that RNA editing can regulate mRNA variant generation. However, the extensive changes in alternative splicing observed upon ADAR1 knockdown could not be completely connected to the editing function, and, indeed, only a few editing sites were detected in the vicinity of near regulatory splicing sequences. Similarly, Licht et al. (2016)^23^ demonstrated that ADAR2 RNA-binding could influence splicing efficiency in an editing-independent manner for specific ADAR2 target minigene constructs. In agreement, Kapoor et al. (2020)^22^ recently showed that Adar −/− mouse-derived tissues display extensive editing-independent splicing changes.

In this work, we investigated the role of ADAR1 in alternative splicing in four transgenic non-related cell lines, focusing on the connections between RNA editing and alternative splicing, showing that most splicing events upon ADAR1 knockdown or overexpression are not related to RNA editing events. We also established the ADAR1p110 protein-protein interactome, showing interactions with several key spliceosome co-factors. Furthermore, using two ADAR1p110 inducible cell lines, we found that most ADAR1-related splicing events do not involve RNA editing near the affected splicing junctions. Finally, we show that overexpression of ADAR1p110 alters the exon usage of splicing factor isoforms within the cell, suggesting that ADAR1 does exert indirect effects on splicing. These results highlight a complex regulatory network of splicing mediated by ADAR1 that includes direct RNA editing and indirect effects through the regulation of the splicing machinery.

## Methods

### Cell Culture

HeLa and HEK293 cells were cultured under standard conditions at 37 °C in a humidified incubator containing 5% CO2. HeLa ADAR1p110 Flp-In T-REx and GFP Flp-In T-REx cells were generated, subcloning the ADAR1p110 cds into the pCDNA5-FRT-cGFP plasmid to further integrate the construct using the Flp-In System following manufacturer instructions (Thermo Fisher Scientific).

### Immunoblotting and co-immunoprecipitation

HeLa ADAR1p110 Flp-In T-REx cells were lysed (one 10 cm dish at 80% (800.000 cells) confluency per immunoprecipitation experiment) in 1mL lysis buffer (10 mM Tris/Cl pH 7.5; 150 mM NaCl; 0.5 mM EDTA; 0.5% Triton X) supplemented with protease inhibitors (Halt, Thermo Fisher Scientific) and 250 units of Benzonase (Sigma Aldrich), and placed on ice for 30 mins. At the same time, extensively shred using a G15 syringe. Cell lysates were centrifuged at 20.000 rpm for 10 min at 4°C, and the supernatant was transferred to a low binding pre-cooled 1.5 mL tube. 40 μL pre-washed control beads (Pierce) (washed three times in cold lysis buffer completed with DTT 1mM and proteinase inhibitors) slurry were added to the lysate and incubated for 30 min at 4°C under rotation. Further, the lysate was spun down at 2500 rpm for 2 min, and the supernatant transferred to a new low binding pre-cooled 1.5 mL and incubated with 40μL of pre-washed GFP-trap beads (washed three times in cold lysis buffer completed with DTT (Thermo Fisher Scientific) 1mM and protease inhibitors) during four h under gentle rotation at 4°C, the complexed beads were spun down at 4°C at 2500 rpm for 2 min and the supernatant was discarded to further wash them three times with lysis buffer (supplemented with protease inhibitors). Finally, the complexed beads were resuspended 40 μL. For co-immunoprecipitation experiments, 10 μL of the complexed beads were boiled with 4x NuPAGE Buffer (Thermo Fisher Scientific) and three μL 1M DTT following manufacturer recommendations. Western blotting was performed as stated previously in Sagredo et al. (2017)^37^, and for mass spectrometry, the beads were eluted by pH elution approach as indicated in the manufacturer’s protocol (GFP-trap_A; Chromotek). Mass spectrometry was performed as previously described in Avolio et al. (2018)^38^.

### Reporter assay and transfections

Reporter assays were performed according to Armisén et al. (2011)^39^. Briefly, HeLa cells were transfected with one µg of the CMV-LUC2CP/ARE plasmid (Addgene #62857), 0.5 µg of Renilla plasmid (pRL-Renilla, Promega), and one µg of the different ADAR1 isoform plasmids. ADAR1p110 deaminase inactive (DeaD) and the ADAR1p110 dsRBD mutant (EAA) were cloned into a pCDNA3.1(+) backbone derived from ADAR1p110 cDNA. Dual-Luciferase measurements were performed using the Dual-Glo Luciferase system kit (Promega), using Cytation 3 Multi-Mode Reader (BioTek Instruments). Renilla signal was used to normalize prior luciferase comparisons between the different samples. Plasmid and siRNA transfections were carried out as described in Sagredo et al. (2020)^18^. Briefly, the different plasmids used in this study were transfected using Lipofectamine 3000, according to the manufacturer’s instructions (Thermo Fisher Scientific), using six-well plates at 80% confluency. For siRNA control (Cell Signaling, 6568S) or ADAR1 (Thermo Fisher Scientific, 119580), siRNAs were transfected at a final concentration of 20 nM and incubated for 48 hours for further experiments. For plasmid transfection, 2.5 µg of the different plasmids were transfected, and 16 h after transfection, the cells were grown with new fresh media to be harvested 48 h after the transfection procedure.

### RNA-seq experiments and workflow

RNA integrity and RNA sequencing were performed as described previously in Sagredo et al. (2020)^18^. Fastq files were aligned using STAR (two-step mode) against hg19, and transcripts were counted with HT-seq software. Differential exon usage (DEU) analysis for the different cell lines and datasets included in this study was performed using the DEXSeq^40^ tool v1.36.0 (DEXseq_count.py script) to perform the differential DEU analysis further as recommended by the software developers. Features with an FDR<0.01 and Log2 > fold one were included in the MDA-MB-231 analysis, and FDR<0.05 and Log2 fold > 0.75 features were included in the ZR-751 analysis. Differential splicing events were called and analyzed using rMATS-turbo v4.1.2 software^41,42^, comparing control samples vs. ADAR1 overexpressing cells. Splice junction definitions from the UCSC goldenPath repository were utilized for further differential splicing analysis. All generated data was processed in R (3.6.1) or the Unix command line. Datasets from ZR-751 and MDA-MB-231 cells are analyzed from Sagredo et al. (2020)^18^ (SRA accession numbers SRP200635 and SRP200634).

### RNA editing and Splicing intersection analysis

Similarly to Kapoor et al. (2020)^22^, we intersected the differential exon usage analyzed using the DEXseq tool. We scanned the called editing sites in 50 nt, 200 nt, 500 nt, and 1000 bp windows on both sides of the exon/intron boundary. In addition, reference sites present in RADAR and DARNED databases were also included^43,44^. This analysis was performed for significant differential exons, included/excluded in the ZR-751 and MDA-MD-231 cells. DARNED and RADAR events were annotated and intersected with the spliceosome iCLIP from Briese et al. (2019)^45^ E-MTAB-8182, using bedtools intersect v2.26.0, and the same window scanning strategy was used to intersect both datasets.

rMATS splicing events were called in default mode (--b1 {bams_controls} --b2 {bams_oe} --gtf hg19.ensGene.gtf -t paired --readLength 100 --nthread 12 –od ./output_files/ --tmp ./output_files/tmp --libType fr-firststrand ). For further analysis, the files counted only the reads that used cross-splice junctions. The splicing events with less than 20 reads between the sum of counts with and without in each replicate, including the exon/intron/splice event, were removed. The Remaining events were filtered by FDR ≤ 0.05, and The remaining events were filtered by FDR ≤ 0.05, with a difference between control and OE groups of | IncLevelDifference | ≥ 0.1. Where IncLevelDifference is the difference of PSI (Percent Spliced In) values of Control vs OE treatment groups. The filtered set of splicing events was then annotated with external_gene_name using the biomaRt package v2.54.0 from Bioconductor. For those significant splicing changes, we selected a set of windows involving the splicing event. The window considered 50 bp from the splice junction into the body of the exons and 200 bp into the introns of the event, except for RI events, where 50 bp was considered for both directions of the splice junctions. This set of windows was created for each significant splicing event and intersected with the set of edited sites. The edited sites were filtered by total depth ≥ 5 (considering all samples, 6 per cell type) and kept those observed at least in 2 replicates or observed in RADAR and/or DARNED databases^43,44^. Then, both sets, splicing events and edited sites, were intersected with bedtools v2.26.0 using bedtools intersect. Also, the splicing events were annotated with RADAR+DARNED, obtained from REDIportal ^46^ using the same “windows set” strategy to account for known editions intersecting the splice site windows. If any splicing event intersects with the sample’s editions or a known edition from RADAR+DARNED, the event is tagged as Editing Dependent; otherwise, it is tagged as Editing-Independent. Finally, rMATS significant changes were additionally annotated and intersected with the ADAR1 iCLIP data from Chen et al. (2015) (GSE63709) ^16^ and Bahn et al. (2015)^14^ (GSE55363) and the irCLASH data from Song et al. (2020)^47^(GSE136327).

### Quantitative RT-PCR

Total RNA was extracted using the Mammalian Total RNA Isolation Kit following the manufacturer’s instructions (SIGMA-Aldrich). This includes Turbo DNAse (Thermo Fisher Scientific) treatment following manufacturer instructions. 500 ng of RNA was reverse-transcribed using an AffinityScript qRT-PCR cDNA Synthesis Kit (Agilent Technologies Inc.) and diluted 4 times. qRT-PCR was performed using specific primers (for further details, please check Supplementary Material 1) and the KAPA SYBR Green master mix (Roche). The reactions were performed using the MIC qPCR machine using the following thermal profile: 95 °C for 15 seconds, 58 °C for 15 seconds, and 72 °C for 15 seconds at 40 cycles, including no template controls. Expression values were calculated using the ΔΔCt method and expressed as the fold of change relative to control samples. ACTNB was used as a housekeeping gene. In addition, RESS-qPCR for AZIN1 and MDM2 targets was performed as described in Crews et al. (2015)^48^. Primers were used to validate DEU data, targeting one specific exon and normalizing it against GAPDH. For rMATS validations, two primers were used for splicing change measurements, amplifying the affected exon and a constant region of the targeted transcript.

### Site level A to G(I) editing comparisons in HeLa and HEK293 FLP T-REx cells

Variant calling files, annotations, and filtering were performed as described previously in Sagredo et al. (2020) ^18^ and Widmark et al. (2022)^49^. Only addressable A to G(I) variants were used, Fisher’s exact test was used to compare the treatments, and a confidence level of 0.05 was used as a cut-off for further analysis. All the data generated was processed in R (3.6.1).

### Gene ontology and pathway enrichment analysis

To determine the relationship between ADAR1 and splicing regulation, a gene ontology analysis was carried out using Cytoscape^50^ v3.7.1 software and ClueGO^51^ v2.5.4 plugin or ShinyGO^52^ 0.76.3 web application. For ClueGO analysis, a gene list was submitted on this software using the Biological Process database (v06/01/2022) for further comparisons. Only statistically significant groups were displayed, using a Bonferroni step-down multiple comparison post-hoc test. A corrected p<0.05 was considered statistically significant. For the protein-protein enrichment analysis, the STRING^53^ database webpage was used, using the complete list (Uniprot IDs) of significant targets (FDR<0.05, compared to GFP samples).

## Results

### Editing of splicing motifs does not correlate with alterations in exon usage upon ADAR1 expression

To investigate ADAR’s involvement in splicing regulation, we performed a differential exon usage analysis datasets generated upon knockdown of ADAR1 by either shRNA (ZR-751 ADAR1 knockdown cells) or overexpression by adenoviral transduction (MDA-MB-231 cells overexpressing ADAR1p110). Our analysis revealed 960 significant exon usage changes in the ZR-751 ADAR1 knock-down cells, compared to controls, and 1958 significant exon usage changes in the MDA-MB-231 ADAR1p110 overexpressing cells, compared to GFP transduced cells, used as control. These changes mapped to 300 and 805 transcripts at the gene level, respectively (Figure 1.A). Those transcripts that exhibited differential exon usage after the alteration of ADAR1 expression were enriched in pathways related to apoptosis, cell mobility, and regulation of cell proliferation (Figure 1.B). Only 24 transcripts overlapped between the datasets, suggesting that ADAR1-dependent splicing regulation strongly varies across cell lines. Interestingly, and despite the low overlap at the gene level, both cell lines showed significant changes in exon usage in transcripts involved in cancer progression, including DUSP6, AKT1, WNT9A, IL6ST, and FGFR1 (Supplementary Table 1).

**Figure 1.**
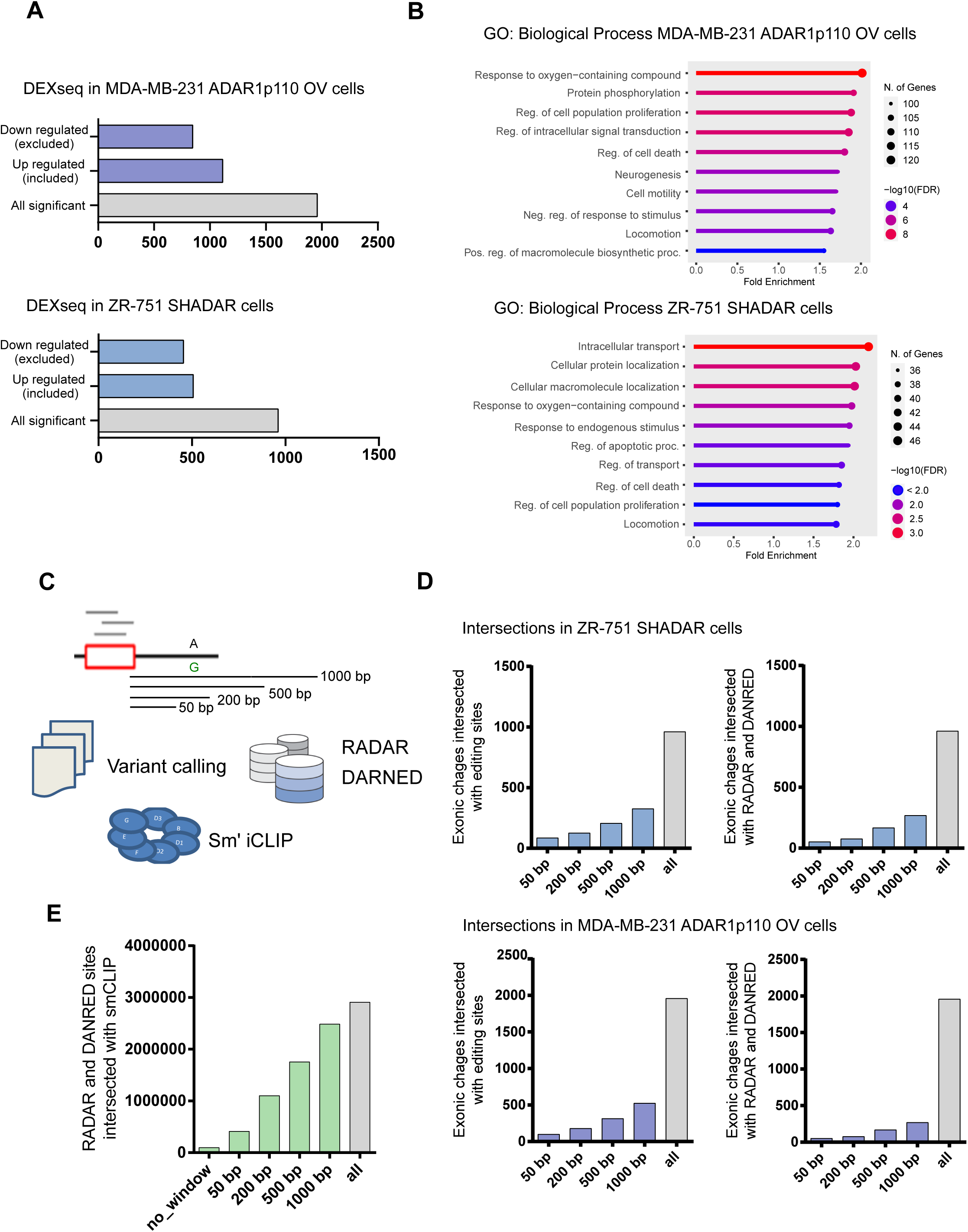
ADAR1 manipulation produces significant splicing changes not entirely associated with their A to I activity. A) Differential Exon Usage (DEU) after ADAR1 manipulation in ZR-751 and MDA-MB-231 cells. DEU analysis comparing MDA-MB-231 ADAR1p110 overexpressing versus GFP transduced cells (upper panel). ZR-751 SHC versus ZR-751 ADAR1 knockdown DEU analysis (lower panel). B) Gene Ontology (Biological process) enrichment analysis for those transcripts with significant DEU in MDA-MB-231 cells (Upper panel) and ZR-751 cells (Lower panel). Showing the significant terms based on gene number and Fold enrichment (FDR). C) Diagram to identify RNA editing events associated with the DEU changes, using the variant calling files, RADAR and DANRED and positional information from Briese et al (2019)^45^, spliceosome CLIP data, using different spanning windows (50 bp to 1 Kbp) to identify possible A to G (I) variants associated to DEU changes found in the MDA-MB-231 and ZR-751 cells, after ADAR1 manipulation. D) Number of DEU with addressable editing sites in the compromised exon position with an extended boundary of 100 bp, 400, 1000, or 2000 bp, using their variant calling information (left panel), RADAR, and DANRED databases (right panel) in ZR-751 cells (upper) and MDA-MB-231 (lower) cells. E) Number of editing sites present in RADAR and DANRED databases and their association with the spliceosome CLIP positions based on Briese et al. (2019)^45^ and their intersection using an extended boundary of 0 bp, 50 bp, 200 bp, 500 bp, or 1000 bp.

To elucidate how ADAR RNA editing is related to splicing regulation, we selected all the differential exons present in both ADAR1 knockdown and ADAR1p110 overexpressing cells and determined if editing sites map to the vicinity of intron boundaries in the compromised mRNAs. To do so, we used the unfiltered RNA variant calling analysis to maximize the detection of any potential RNA editing site. This data was intersected with the coordinates of compromised exons applying windows ranging from 50 to 1000 base pairs (bp) in either direction. We also considered all the editing sites annotated in the RADAR and DARNED databases to include any previously described editing sites that may not have been identified in the knockdown and overexpression cells (Figure 1.C). As expected, the overlap between differentially used splice sites and edited bases increased as the window size increased. Despite applying the bigger window (1000 bp), only ∼30% of the differentially used exons overlapped with editing sites. This lack of overlap was observed when RADAR and DARNED editing sites were used (Figure 1.D). These surprising results suggest that most alternative exon usage events are associated with ADAR-independent editing, even when using very permissive parameters.

To reinforce these results, we compiled the spliceosome binding sites reported by previous iCLIP-published datasets. We tested the binding footprints on the RNA against RNA editing events collected above. The overlapping between the spliceosome iCLIP and the editing databases RADAR and DANRED was only 3.33%, yielding 96896 editing sites from the 2909674 total sites in RADAR and DANRED. However, the overlapping increased to 85.4% when a 1000 bp window was used. Altogether, these complementary analyses revealed an apparent lack of correlation between ADAR1-mediated RNA edits and alternative splicing events (Figure 1.E).

### The protein-protein interactome of ADAR1p110 includes many splicing factors

Classical biochemistry assays have associated ADAR1 with RNA-binding proteins involved in splicing, such as HNRNPL, NONO, and SFPQ^54^. To study all the protein partners of ADAR1p110, we generated HeLa Flp-In Trex cells that express a GFP-tagged ADAR1p110 protein in a doxycycline-dependent manner (Supplementary Figure 1). ADARp110 was immunoprecipitated using GFP-trap beads under stringent conditions, and interactors were identified by label-free quantitative proteomics (Figure 2A). GFP-expressing control cells were used to exclude contaminants. The analysis revealed 432 significantly enriched in ADAR1p110-GFP. As immunoprecipitation was performed in native conditions, these proteins reflect direct and indirect protein-bridged interactors. Notably, 270 of the interactors are annotated as RBPs, 163 of which harbor classical RNA-binding domains (RBDs), and 107 are non-canonical^55,56^ (Figure 2B and Supplementary Table 2). To focus on the most prominent (stoichiometric) interactors, we selected the proteins with a higher eluate signal (q50 quantile of the IBAQ signal). We performed a pathway enrichment analysis using Reactome and STRING databases. We found a significant enrichment of proteins involved in ‘ribosome’ (FDR: 3.41E-80), ‘spliceosome’ (FDR: 1.49E-38), ‘RNA transport’ (FDR: 0.0273), ‘mRNA surveillance’ (FDR: 4.21E-23) and ‘mismatch repair’ (FDR: 9.53E-17). (Table 2. C and Supplementary Table 3). We observed a striking enrichment of members of the Sm complex and U2 snRNP complex in ADARp110 interactome (SF3A3, SF3B2, SF3B3, SNRPB2, SNRPA1, SNRPD3, SNRPE and SNRPG)^57^ (Supplementary Table 2). We also identified splicing regulators such as HNRNPL, DHX15, SFPQ, and STRBP^58^. We validated several of the observed interactions by ADAR1 immunoprecipitation followed by western blot showing that Sm antigen, the splicing factor SF3A3, and two different heteronuclear proteins, HNRNPL and HNRPA/B, reinforcing the significance of our results (Figure 2D). These results imply that ADAR1 may also regulate splicing through interaction with the spliceosome and several of its cofactors.

**Figure 2.**
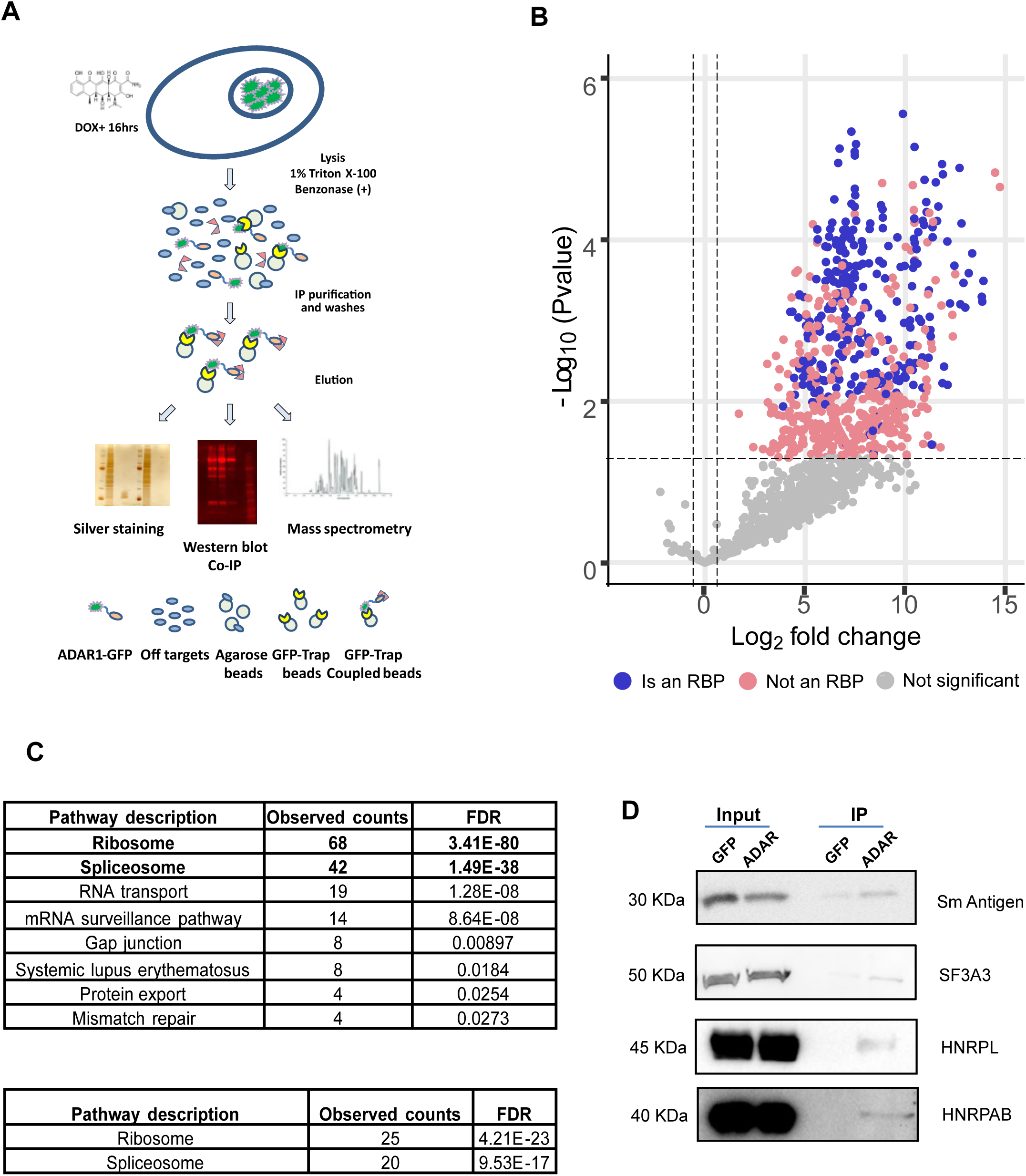
HR-MS reveals that ADAR1p110 interacts with splicing machinery. A) Model of work for HR-MS to capture ADAR1-p110 protein interactors. B) Volcano plots showing significant ADAR protein-protein interactors, highlighting significant (pink) and known RNA binding proteins (RBP) (blue) interactors. Dashed lines depict significant -Log10 p-value (FDR) (y-axis> 1.75) and Log2 Fold change over GFP (x-axis> 1). C) Reactome/STRING pathway enrichment for Q1 and Q2 IBAQ ranked ADAR1p110 interaction partners. D) co-IP validations in HeLa cells overexpressing GFP or ADAR1p110, showing inputs and IP elutions for Sm Antigen, SF3A3, HNRPL, and HNRPAB.

### Deconvoluting the role of ADAR1p110 in splicing

Our results indicate an editing-independent function of ADAR1 in splicing, consistent with many splicing factors in ADAR’s interactome. We induced the expression of ADAR1p110-GFP in HeLa and HEK293 Flp-in TREx cells and analyzed the exon and splicing changes after ADAR1p110 induction. Using DEXseq, we observed that 4207 and 4235 exons had decreased usage in HeLa and HEK293, while 3751 and 2720 exons showed higher usage than GFP-induced control cells (Fig 3A and Supplementary Table 4). To characterize the transcripts with changes in exon usage, we compared the results in both lines at the transcript level, showing 1712 transcripts differentially spliced in an ADAR-dependent manner in both cell lines. These genes are involved in ‘RNA processing’ (FDR: 1.4E-30), ‘protein localization’ (Protein localization to organelle (FDR: 4.8E-24), and ‘viral processing’ (FDR: 3.2E-22) GO terms, among others. These terms are highly enriched in RBPs and splicing-related factors (Figure 3. B and Supplementary Table 5).

**Figure 3.**
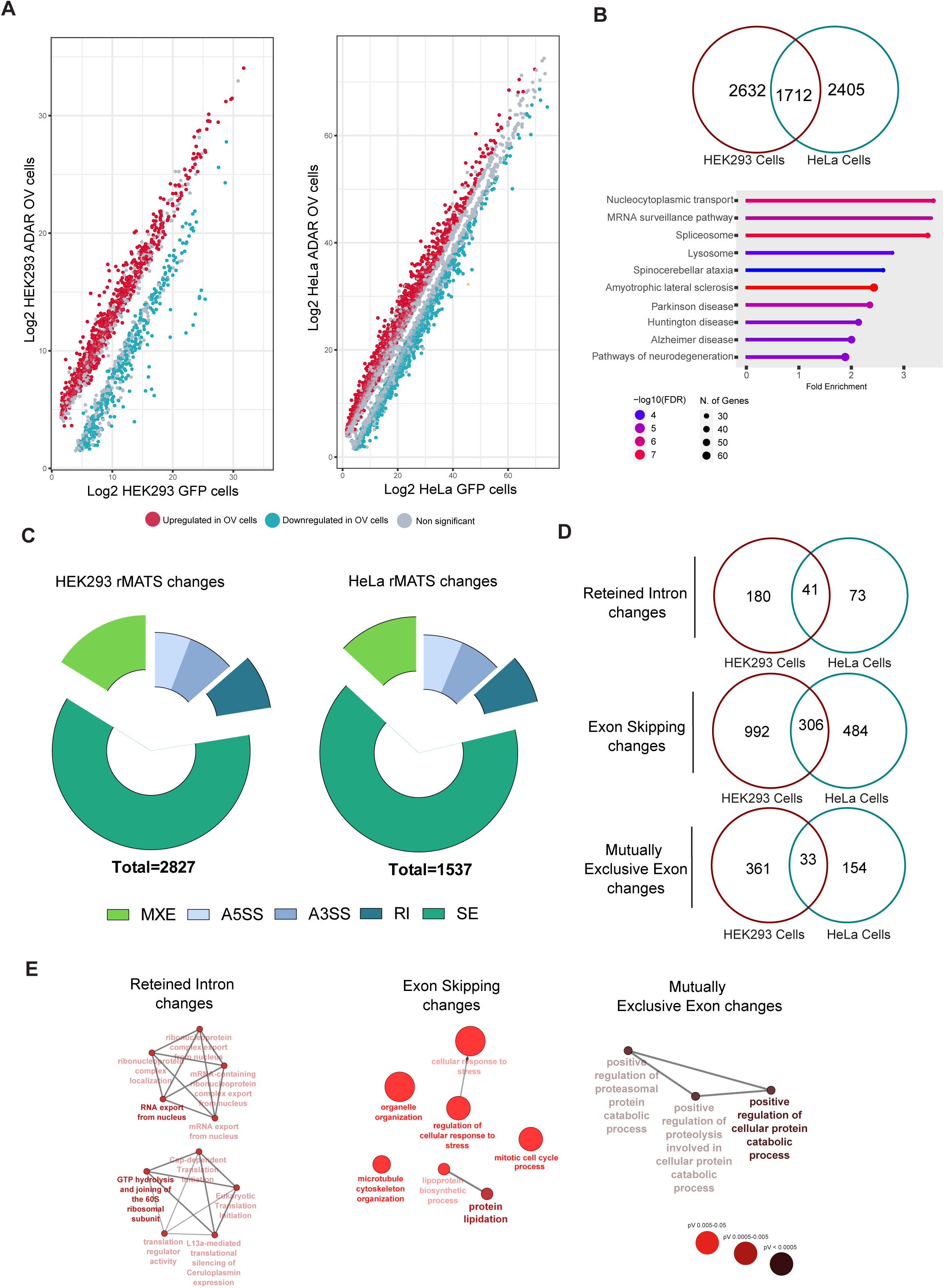
ADAR overexpression on HeLa and HEK293 models produces significant splicing changes. A) Differential exon usage counts (DEU) on HEK293 (left) and HeLa ADAR1p110 overexpressing cells, showing those exons significantly overrepresented (included) in red and those excluded after ADAR1p110 overexpression (in green). B) The Venn diagram shows the overlap -at transcript level-between those significant DEU changes in HEK293 and HeLa cells (upper), showing the significant biological processes associated with those shared transcripts with significant DEU in both cell lines. C) Pie charts showing the significant splicing change types after rMATS analysis in HEK293 ADAR1p110 (left) and HeLa ADAR1p110 (right) overexpressing cells. D) Venn diagrams showing the overlap in the different splicing event changes between HEK293 and HeLa ADAR1p110 overexpressing cells. E) Gene enrichment analysis (Biological process) for those splicing events shared between HEK293 and HeLa ADAR1p110 cells.

Further, we utilized rMATS to analyze splicing events, an event-based software that circumvents the limitations of differential exon usage based on exonic counts. Using this tool, we detected 1537 significant alternative splicing changes in the HeLa cells and 2827 in the HEK293 cells upon ADAR1p110-GFP overexpression, compared to GFP-induced cells (Figure 3C). Most of the changes were related to exon skipping (SE), representing 65,7% and 61,4% in HeLa and HEK293 cells, respectively, upon ADAR1p110 overexpression. The following most abundant splicing events were mutually excluded exon changes (MXE) (13% and 16.1%, respectively), intron retention (RI) (7.9% and 8.9%, respectively), and alternative 5’ and 3’ splice site usage (13.3% and 13.6%, respectively) (Figure 3D and Supplementary Table 6 and Supplementary Table 7), indicating that ADAR1p110 exerts alternative splicing changes on many transcripts across different cell types. However, many of the observed transcripts that exhibited splicing changes do not overlap between cell lines, suggesting that regulation of alternative splicing by ADAR1-p110 depends on the cell in which it is expressed (Fig 3D).

GO analysis revealed that transcripts with intron retention changes in both cell lines are enriched in RNA splicing and RNA processing (FDR: 0.042) and Cytoplasmic translation (FDR:0.042). The transcripts that exhibited significant exon skipping changes are enriched in ‘protein lipidation,’ ‘organelle organization’ (Organelle assembly and microtubule cytoskeleton organization, FDR: 0.009), and DNA damage repair related GO terms (i.e., ‘mitotic cell cycle’ FDR: 0.009). Conversely, significant mutually exclusive exon changes were enriched in protein catabolic-related processes (‘pos. reg of cellular protein metabolic processes’, FDR: 0.02) (Figure 3.E). Finally, we found that several splicing factors, such as THOC1, ACIN1, TIA1, SRSF6, HNRNPD, DDX46, and U2AF1, exhibited significant alterations to their splicing patterns upon ADAR1p110 induction. Furthermore, genes with central roles in the cell cycle and DNA damage response, such as BRCA1, PAX6, MCM3, MKI67, and BCLAF1, also displayed differential splicing in both cell lines.

Our next goal was to validate the ADAR1p110-induced changes in alternative splicing, testing their dependence on RNA editing and, on the RNA-binding activity of ADAR1p110. To do so, we transfected HEK293 cells with an editing deficient ADAR1 mutant (ADAR1-DeaD) and an RNA-binding deficient mutant (ADAR1-EAA) (Supplementary Figure 2.A). We tested the alternative splicing of mRNAs altered in our dataset, including PTPR, HNRNPC, LIG1, NME, and SRSF1. Using splice-site-specific primers, we tested exon exclusion and inclusion. Our results revealed that ADAR1 WT and its mutants induce similar changes in splicing, which was in line with our sequencing analyses. These results suggest that ADAR1p110 can affect the splicing in an RNA-binding and activity-independent manner (Supplementary Figure 2).

### Elucidating the editing dependence of ADAR1-mediated changes to differential splicing

To characterize the editing-independent role of ADAR1p110 in alternative splicing, we performed an A-to-G variant calling on the whole transcriptome of HeLa and HEK293 T-REx cells and intersecting those sites with the altered splicing junctions in our data. As expected, most of the detected editing sites (∼90%) occurred in Alu-elements situated in non-coding genomic regions (Supplementary Table 4), and less than 0.3% of detected edited sites were found in coding regions (Figure 4A). In addition, we performed a GO analysis on the significant edited sites shared between HEK293 and HeLa cells at the gene level (1598 genes), showing enrichment for ‘cell cycle’ (FDR:0.0001), ‘DNA replication’ (FDR:0.002), ’Fanconi anemia pathway’ (FDR: 0.001), ‘mismatch repair’ (FDR:0.04), and ‘homologous recombination’ (FDR:0.02) GO terms, among others (Figure 4. B and Supplementary Table 5).

**Figure 4.**
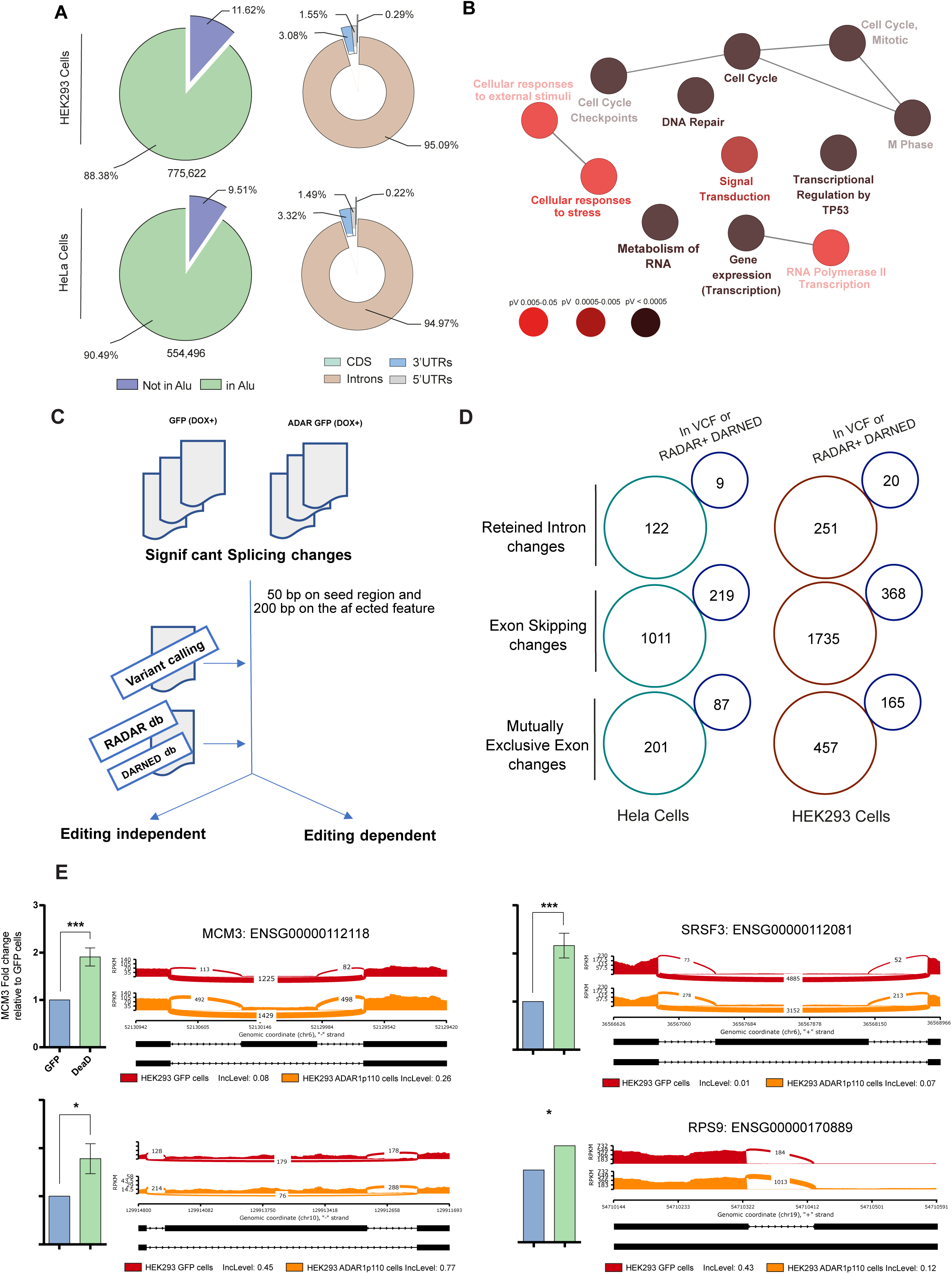
ADAR1p110 overexpression on HeLa and HEK293 cells produces significant splicing changes that are not entirely associated with their A to I activity. A) RNA editing characterization in HeLa and HEK293 cells after performing variant calling, showing the number of editing sites, their distribution on Alu regions and across the mRNA features b) Differentially RNA editing analysis for those shared targets significantly edited in HeLa ADAR1p110 and HEK293 ADAR1p110, compared to GFP overexpressing cells, showing significant Reactome gene ontologies. C) Workflow to identify and recall splicing variants that are affected by editing events and those recalled as editing independent events, briefly rMATS significant sites were used to intersect further the variant calling files and RADAR+DANRED databases, for the intersections of two regions that compromise 250 pb were used (for further details, see material and methods). D) Identification of RNA editing-dependent and RNA-independent events on HeLa and HEK293 cells, showing the different splicing change types on the data and the number of events (in blue) that can be described as editing-dependent. E) qRT-PCR validation for rMATS splicing events recalled as editing independent events after overexpressing HEK293 cells with an empty vector, ADAR1p110, and the DeaD ADAR1p110 plasmid. In addition, sashimi plots are shown for those targets that qRT-PCR validated. Two-tailed student’s T-test was used to calculate differences between samples in E (n=3) (*=P<0.05, **=P<0.01, ***=P<0.001, ****=P<0.0001).

We designed a workflow to intersect the differential splicing events with the RNA editing variants in our data and those RNA editing sites included in different RNA editing databases, considering the splice junction and the intronic region nearby of any significant splicing change (Figure 4. C and Materials and Methods). Interestingly, less than 5% of the splicing changes in our data intersected with RNA editing variants called in the cell lines (Supplementary Figure 3. A and 3. B, right panels), suggesting that most changes cannot directly relate to RNA editing. From the different splicing changes, mutually exclusive exon (MXE) changes had the highest number of edited sites in their sequence. In contrast, the 5’ splice site, 3’ splice site, and intron retention changes had the lower frequency of RNA editing sites in their coordinates (Figure 4D, Supplementary Table 6 and Supplementary Table 7), where a little fraction intersected in RADAR, DARNED and/or Reditools databases (Figure 4.D, Supplementary Table 7 and Supplementary Figure 3. A and 3.B). Further, we selected several transcripts that exhibited alternative splicing changes in the absence of RNA editing for validation by qRT-PCR. Those targets are related to DNA damage response and alternative splicing, including MCM3, RPS9, MKI67, and SRSF3. Strikingly, we observed that overexpression of ADAR1-DeaD mutant led to significant changes in the alternative splicing of these transcripts, supporting that the alteration of their splicing is editing-independent (Figure 4E).

### ADAR1p110 regulates and contributes to transcriptome variability affecting mRNAs that encode for splicing cofactors

Our results showed that many ADAR1-regulated alternative splicing appears to be editing-independent significantly. To test this phenomenon, we co-transfected the different ADAR1p110 mutants with a splicing reporter consisting of a luciferase split with an intron. Active luciferase synthesis thus requires splicing. We observed a notable decrease in luciferase signal upon ADAR1p110-DeaD and ADAR1p110-EAA overexpression in HEK293 cells (Figure 5.A). These results agree with Kapoor et al. (2020)^22^ and suggest that non-catalytically active ADAR1p110 influences the splicing of the reporter in both an editing-independent and RNA-binding-independent manner. We had two hypotheses: 1) ADAR regulates splicing factor function through direct protein-protein interactions by changing their RNA-binding properties or stability; 2) the activity of ADAR on mRNAs induces changes in splicing that alter the configuration of splicing factors, leading to differential effects.

**Figure 5.**
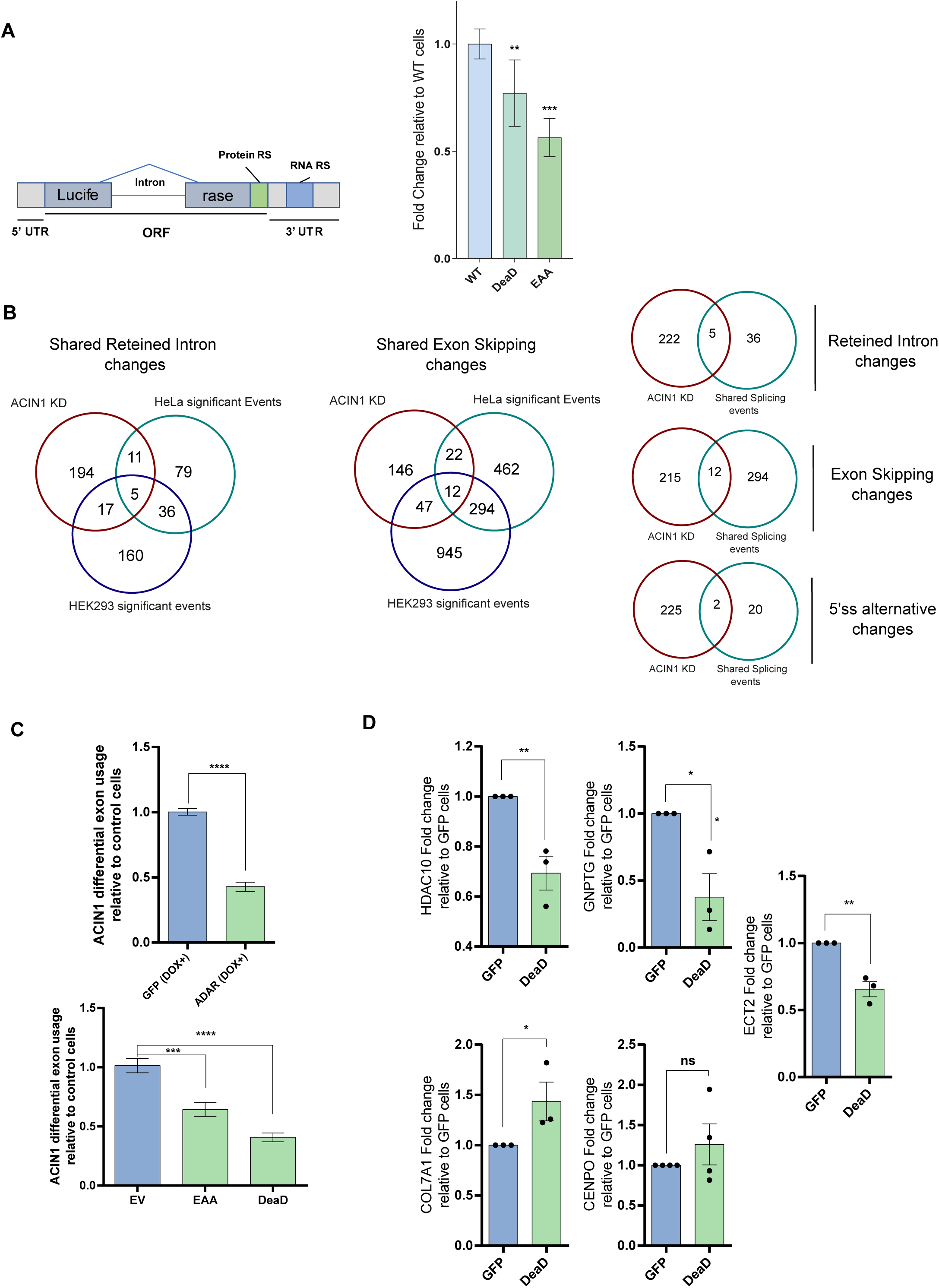
ADAR manipulation produces essential changes in splicing activity and the alternative splicing of other splicing factors. A) Splicing reporter activity after ADAR1 mutant overexpression. Briefly, a split Luciferase plasmid was co-transfected with the different ADAR1p110 mutants, and data was normalized using a renilla plasmid. B) Intersection between splicing events produced by ACIN1, after ACIN1 knockdown and the significant splicing events found in either HeLa or HEK293 ADAR1p110 overexpressing cells, displaying the intersection for all the different splicing event types. C) ADAR overexpression leads to a significant change in the splicing pattern of ACIN1 independent of the editing activity. D) qRT-PCR measuring ACIN1 splicing events after ADAR1 DeaD overexpression, showing significant changes in the splicing pattern of previously validated ACIN1 targets. Two-tailed T-test was used to calculate significant differences between samples (*=P<0.05, **=P<0.01, ***=P<0.001, ****=P<0.0001) in A, C, and D (n=3).

One splicing co-factor affected by ADAR overexpression was ACIN1, a protein part of the apoptosis and splicing-associated protein complex (ASAP) that interacts with the exon junction complex (EJC)^59,60^. The transcript of ACIN1 consistently showed editing-independent alternative splicing and expression changes in both our HeLa and HEK293 ADAR1p110 overexpressing cells (Figure 5.C). Indeed, the overexpression of both DeaD and EAA ADARp110 in HEK293 cells showed a significant change in ACIN1 isoform pattern, both at mRNA and protein levels (Supplementary Figure 2.B), suggesting that ADAR1 could indirectly modify the splicing process by affecting the splicing configuration, result that we expanded to other splicing factors (Supplementary Figure 2.C). To evaluate if the second hypothesis is correct, we intersected publicly available transcriptome data from studies investigating the effect of alternative splicing after ACIN1 knockdown^60,61^, with our data produced after overexpression of ADAR1p110 (Figure 5.B). Interestingly, some of the targets were shared and affected the same mRNA splicing variants between the ACIN1 dataset and our experiments. Further, we performed qRT-PCR in the same splicing regions associated with ACIN1 using the DeaD ADAR1p110 variant. Notably, the DeaD ADAR1p110 overexpression can partially reassemble the effects of ACIN1 (Figure 5.D). Together, these results reinforce the idea that ADAR1 can indirectly regulate alternative splicing of specific transcripts through modulation of alternative splicing on transcripts that code for splicing factors, revealing an intricate role of ADAR1 in regulating alternative splicing and downstream phenotypes.

## Discussion

In this work, we provided new evidence supporting the role of ADAR1p110 in regulating alternative splicing. It has previously been suggested that ADAR1 interacts with the spliceosome and that ADAR1p110 can be detected in the same fraction as U1 and U2 spliceosome complexes^62^. Previous reports have shown that the editing function can be regulated by splicing co-factors such as SRSF9 and DDX15^63,64^ and that RNA immunoprecipitation experiments for HNRNP proteins interact with Alu regions and other features strongly associated with ADAR1 binding sites^54,65^. Moreover, some cofactors of the spliceosome, such as U2AF2, can be detected near editing sites^66^, suggesting a plausible idea that these factors could interact with or displace ADAR1, modifying ADAR1 activity. Our ADAR1p110 protein-protein interactome experiment showed that many ADAR1p110 interactors are indeed RNA binding proteins involved in RNA metabolism and alternative splicing processes, which agrees with other recent ADAR1 proteomic works^67,68^, strengthening the argument that ADAR1p110 could act as a splicing regulator.

We investigated the changes in alternative splicing transcriptome-wide following manipulation of ADAR1 expression and found thousands of changes in each of the four individual cell lines tested. We observed that few of the detected differential splicing events are shared between the two cancer cell lines (ZR-751 and MDA-MB-231), and a significant proportion of those changes present in the HEK293 and HeLa ADARp110-GFP cell lines generated by us, where most of the observed splicing events appear to be cell line specific. This suggests that the regulation imposed by ADAR1 on alternative splicing depends on the particular subcellular environment, confirming the data of previous reports^22^. Furthermore, our data indicates that more than half of the observed differential exon usage in ZR-751 and MDA-MB-231 ADAR1 manipulated cells do not harbor an editing site nearby. Even when using a screening window of 1000 bp and relaxed criteria for editing determination, we found no correlation between splice sites and editing in agreement with previous work^69^.

Upon ADAR1p110-DeaD overexpression, we noted significant changes in the luciferase reporter plasmid, noting that the overexpression of ADAR1p110-EAA also inhibited the luciferase reporter’s splicing when compared to the WT protein. In agreement with this, previous works^21,22^ also investigated the action of an editing-deficient and RNA-binding deficient ADAR1 enzyme and observed that expression of an RNA-binding deficient *Adar1* protein exhibited a more substantial impact on the splicing landscape when compared to the editing-deficient *Adar1* in mice, suggesting RNA editing independent effects. In further support of the editing-independent regulation of alternative splicing by ADAR1, we observed a minor overlap of known edited sites when intersecting their positions with the published iCLIP data for the spliceosome^45^, suggesting an alternative way that the editing-independent functions of ADAR1 are responsible for the majority of the observed differential splicing following its dysregulation.

After overexpressing ADAR1p110-GFP in HEK293 and HeLa cells, we observed that the differentially spliced transcripts were enriched for DNA repair, mRNA surveillance pathways, Splicing/spliceosome, and other related GO terms. Interestingly, many of the observed changes to differential splicing generated transcripts annotated as coding for novel protein isoforms. This phenomenon was previously demonstrated for the ADAR-regulated differential splicing of the CCD15 and RELL2 transcripts^24^. Although the extent of the ADAR1p110 mediated perturbations to alternative splicing remains to be investigated, it may influence mRNA decay or protein isoforms, favoring mRNA variants that are targeted for degradation to mechanisms such as NMD or producing novel protein isoforms. These results could also add an extra layer of complexity in the pathological contexts linked to ADAR1, where this protein is significantly overexpressed and could influence transcript stability and isoform generation, alterations implicated in many disorders and types of cancers^69,70^.

ADAR1p110-GFP overexpression affected the splicing and expression of ACIN1, a core component of the ASAP complex involved in alternative splicing^59^, nonsense-mediated decay^61^, apoptosis, and chromatin condensation^71^. After intersecting the observed differential splicing data in our HEK293 and HeLa cells with differential splicing data available for ACIN1 knockdown^61^, we could determine that a certain proportion of splicing events exhibited overlap between the different datasets, showing the same type of differential splicing on sharing gene coordinates. This suggests that the observed perturbations in splicing could be due to the effect that ADAR1p110 exerts on ACIN1 and may not be due to any direct relationship between ADAR1p110 on those transcripts, results that we further validated by qRT-PCR.

RNA editing occurs co-transcriptionally, and most sites are present in intronic regions^30,72^; a critical challenge of these studies to correlate editing with alternative splicing is enough read coverage to limit false positive events. As our RNA-seq data is based on a poly-A enriched strategy, which limits intronic regions, we supplemented our editing data with the extensive data contained in Reditools, RADAR, and DARNED databases to increase the sensitivity with capturing splicing events that could be associated with RNA editing. However, even with very relaxed parameters (1000 bp) in each direction from the differentially used exons, less than 30% contained edited sites near splicing junctions. Seeing that ADAR1p110 also affects the splicing of multiple other splicing factors and interacts with them at a protein-protein level, the full extent of ADAR1’s involvement in alternative splicing should be investigated systematically to determine these effects in cis or trans.

---

**Supplementary Figure 1. Characterization of the ADAR1p110 Flp-In TREx system.** A) schema of Flp-In TREx ADAR1p110 GFP doxycycline-inducible system. A) GFP fluorescence intensity signal produced in HeLa ADAR1p110-GFP cells after doxycycline induction, recorded during 24h. C) mRNA levels of ADAR1 after doxycycline induction on HEK and HeLa cells. D) RESS-qPCR for RNA editing targets (AZIN1 and MDM2) on HeLa and HEK293 ADAR1p110 after and before doxycycline induction. D) Western blot showing ADAR1p110-GFP after and before doxycycline induction, probing against GFP (upper panel); and confocal imaging showing ADAR1p110 localization, showing nuclear staining (DAPI) and ADAR1p110-GFP (FITC) (63X oil magnification) Two-tailed student’s T-test was used to calculate differences between samples in C (n=3) (**=P<0.01, ***=P<0.001, ns: non-significant).

**Supplementary Figure 2. ADAR1 manipulation produces significant exon usage changes on transcripts.** A) ADAR mutants used for the different transfections, showing the different mutations present across ADAR1p110 domains. B) qPCR validations for differential exon usage not related to ADAR1 editing activity, showing significant changes after DeaD and EAA overexpression in HEK293 cells for PTPRs, HNRNPC, LIG1, NME and SRSF1. C) Representative Western blot showing ACIN1 isoforms after ADAR1 knockdown (siRNA), ADAR1p110 overexpression, and ADAR1 DeaD overexpression (left) in HEK293 cells, with their respective quantification for those bands associated with the larger isoform (200 KDa, 150 KDa, and 90 KDa) (right). A two-tailed T-test was used to calculate significant differences between samples (ns= non-significant, *=P<0.05, **=P<0.01, ***=P<0.001) in B and C (n=3).

**Supplementary Table 1.** Gene enrichment for those common transcripts affected after ADAR1 manipulation in MDA-MB-231 and ZR-751 cells.

**Supplementary Table 2.** ADAR1p110-GFP protein-protein interactome hits, showing the different proteins associated with ADAR1p110 compared to GFP.

**Supplementary Table 3.** STRING and Reactome enrichment analysis for ADAR1p110-GFP protein-protein interactome.

**Supplementary Table 4.** Differential exon usage changes in HeLa and HEK293 cells after ADAR1p110 induction.

**Supplementary Table 5.** GO (BP) enrichment analysis for those transcripts with DEU changes in HeLa and HEK293 cells after ADAR1p110 induction.

**Supplementary Table 6.** rMATS detected changes in HeLa cells after ADAR1p110 induction, annotated using RADAR and DARNED, Variant calling, spliceosome iCLIP, and ADAR1 iCLIP available data.

**Supplementary Table 7.** rMATS detected changes in HEK293 cells after ADAR1p110 induction, annotated using RADAR and DARNED, Variant calling, spliceosome iCLIP, and ADAR1 iCLIP available data.

## Data availability

The data associated with this article are available in its online supplementary material. Raw data will be made available on request.

## Acknowledgments

We thank Dr. Albin Widmark and Dr. Yanara A. Bernal for their feedback on the project and critical reading of the article. We acknowledge support from the National Genomics Infrastructure in Stockholm funded by Science for Life Laboratory, the Knut and Alice Wallenberg Foundation and the Swedish Research Council, and the Swedish National Infrastructure for Computing/Uppsala Multidisciplinary Center for Advanced Computational Science for assistance with massively parallel sequencing and access to the UPPMAX computational infrastructure.

## Author contributions

Conceptualization, E.A.S., R.A. and A.B.; investigation, E.A.S., V.K., P.M., M.V., A.B., A.I.J. and M.V., visualization, E.A.S., A.B., writing-review and editing, E.A.S., V.K., K.M., A.C. and R.A. supervision, E.A.S., N.V., R.A., project administration and funding acquisition, R.A., N.V. and E.A.S. All authors have read and agreed to the published version of the manuscript.

## References

1. Nishikura, K. Functions and regulation of RNA editing by ADAR deaminases. Annual Review of Biochemistry 79, 321–349 (2010).

2. Tan, M. H. et al. Dynamic landscape and regulation of RNA editing in mammals. Nature 550, 249–254 (2017).

3. Bass, B. L. RNA editing by adenosine deaminases that act on RNA. Annual Review of Biochemistry 71, 817–846 (2002).

4. Keegan, L. P., Gallo, A. & O’Connell, M. A. The many roles of an RNA editor. Nature Reviews Genetics 2, 869–878 (2001).

5. Ramaswami, G. et al. Identifying RNA editing sites using RNA sequencing data alone. Nature Methods 10, 128–132 (2013).

6. Peng, Z. et al. Comprehensive analysis of RNA-Seq data reveals extensive RNA editing in a human transcriptome. Nature Biotechnology 30, 253–260 (2012).

7. Savva, Y. A., Rieder, L. E. & Reenan, R. A. The ADAR protein family. Genome biology vol. 13 252 at 10.1186/gb-2012-13-12-252 (2012).

8. Eckmann, C. R., Neunteufl, A., Pfaffstetter, L. & Jantsch, M. F. The human but not the Xenopus RNA-editing enzyme ADAR1 has an atypical nuclear localization signal and displays the characteristics of a shuttling protein. Molecular Biology of the Cell (2001) doi:10.1091/mbc.12.7.1911.

9. Poulsen, H., Nilsson, J., Damgaard, C. K., Egebjerg, J. & Kjems, J. CRM1 Mediates the Export of ADAR1 through a Nuclear Export Signal within the Z-DNA Binding Domain. Molecular and Cellular Biology (2001) doi:10.1128/mcb.21.22.7862-7871.2001.

10. Hideyama, T. et al. Profound downregulation of the RNA editing enzyme ADAR2 in ALS spinal motor neurons. Neurobiology of Disease (2012) doi:10.1016/j.nbd.2011.12.033.

11. Laxminarayana, D., O’Rourke, K. S., Maas, S. & Olorenshaw, I. Altered editing in RNA editing adenosine deaminase ADAR2 gene transcripts of systemic lupus erythematosus T lymphocytes. Immunology 121, 359–369 (2007).

12. Han, L. et al. The Genomic Landscape and Clinical Relevance of A-to-I RNA Editing in Human Cancers. Cancer Cell 28, 515–528 (2015).

13. Solomon, O. et al. Global regulation of alternative splicing by adenosine deaminase acting on RNA (ADAR). Rna 19, 591–604 (2013).

14. Bahn, J. H. et al. Genomic analysis of ADAR1 binding and its involvement in multiple RNA processing pathways. Nature Communications 6, 6355 (2015).

15. Ota, H. et al. ADAR1 forms a complex with dicer to promote MicroRNA processing and RNA-induced gene silencing. Cell 153, 575–589 (2013).

16. Chen, T. et al. ADAR1 is required for differentiation and neural induction by regulating microRNA processing in a catalytically independent manner. Cell Research 25, 459–476 (2015).

17. Wang, I. X. et al. ADAR Regulates RNA Editing, Transcript Stability, and Gene Expression. Cell Reports 5, 849–860 (2013).

18. Sagredo, E. A. et al. ADAR1 Transcriptome editing promotes breast cancer progression through the regulation of cell cycle and DNA damage response. Biochimica et Biophysica Acta - Molecular Cell Research 1867, (2020).

19. Karlström, V. et al. ADAR3 modulates neuronal differentiation and regulates mRNA stability and translation. Nucleic Acids Research gkae753 (2024) doi:10.1093/nar/gkae753.

20. Licht, K. & Jantsch, M. F. The Other Face of an Editor: ADAR1 Functions in Editing-Independent Ways. BioEssays vol. 39 at 10.1002/bies.201700129 (2017).

21. Licht, K., Kapoor, U., Mayrhofer, E. & Jantsch, M. F. Adenosine to Inosine editing frequency controlled by splicing efficiency. Nucleic Acids Research 44, 6398–6408 (2016).

22. Kapoor, U. et al. ADAR-deficiency perturbs the global splicing landscape in mouse tissues. Genome Research 30, 1107–1118 (2020).

23. Licht, K. et al. A high resolution A-to-I editing map in the mouse identifies editing events controlled by pre-mRNA splicing. Genome Research 29, 1453– 1463 (2019).

24. Tang, S. J. et al. Cis- and trans-regulations of pre-mRNA splicing by RNA editing enzymes influence cancer development. Nature Communications 11, (2020).

25. Egebjerg, J., Kukekov, V. & Heinemann, S. F. Intron sequence directs RNA editing of the glutamate receptor subunit GluR2 coding sequence. Proceedings of the National Academy of Sciences of the United States of America 91, 10270–10274 (1994).

26. Schoft, V. K., Schopoff, S. & Jantsch, M. F. Regulation of glutamate receptor B pre-mRNA splicing by RNA editing. Nucleic Acids Research 35, 3723–3732 (2007).

27. Kawahara, Y., Ito, K., Ito, M., Tsuji, S. & Kwak, S. Novel splice variants of human ADAR2 mRNA: Skipping of the exon encoding the dsRNA-binding domains, and multiple C-terminal splice sites. Gene (2005) doi:10.1016/j.gene.2005.07.028.

28. 28. Rosenthal, J. J. C. & Seeburg, P. H. A-to-I RNA Editing: Effects on Proteins Key to Neural Excitability. Neuron at 10.1016/j.neuron.2012.04.010 (2012).

29. Rueter, S. M., Dawson, T. R. & Emeson, R. B. Regulation of alternative splicing by RNA editing. Nature 399, 75–80 (1999).

30. Hsiao, Y. H. E. et al. RNA editing in nascent RNA affects pre-mRNA splicing. Genome Research 28, 812–823 (2018).

31. Nilsen, T. W. & Graveley, B. R. Expansion of the eukaryotic proteome by alternative splicing. Nature vol. 463 457–463 at 10.1038/nature08909 (2010).

32. Hartner, J. C. et al. Liver Disintegration in the Mouse Embryo Caused by Deficiency in the RNA-editing Enzyme ADAR1. Journal of Biological Chemistry 279, 4894–4902 (2004).

33. Hartner, J. C., Walkley, C. R., Lu, J. & Orkin, S. H. ADAR1 is essential for the maintenance of hematopoiesis and suppression of interferon signaling. Nature Immunology 10, 109–115 (2009).

34. XuFeng, R. et al. ADAR1 is required for hematopoietic progenitor cell survival via RNA editing. Proceedings of the National Academy of Sciences of the United States of America 106, 17763–17768 (2009).

35. Bajad, P. et al. An internal deletion of ADAR rescued by MAVS deficiency leads to a minute phenotype. Nucleic Acids Research 48, 3286–3303 (2020).

36. Heraud-Farlow, J. E. et al. Protein recoding by ADAR1-mediated RNA editing is not essential for normal development and homeostasis. Genome Biology 18, (2017).

37. Sagredo, A. I. et al. TRPM4 regulates Akt/GSK3-β activity and enhances β-catenin signaling and cell proliferation in prostate cancer cells. Molecular Oncology 12, 151–165 (2018).

38. Avolio, R. et al. Protein Syndesmos is a novel RNA-binding protein that regulates primary cilia formation. Nucleic Acids Research 46, 12067–12086 (2018).

39. Armisén, R. et al. TRPM4 enhances cell proliferation through up-regulation of the β-catenin signaling pathway. Journal of Cellular Physiology 226, 103–109 (2011).

40. Anders, S., Reyes, A. & Huber, W. Detecting differential usage of exons from RNA-seq data. Genome Research 22, 2008–2017 (2012).

41. Shen, S. et al. rMATS: Robust and flexible detection of differential alternative splicing from replicate RNA-Seq data. Proceedings of the National Academy of Sciences of the United States of America 111, E5593–E5601 (2014).

42. Wang, Y. et al. rMATS-turbo: an efficient and flexible computational tool for alternative splicing analysis of large-scale RNA-seq data. Nature Protocols (2024) doi:10.1038/s41596-023-00944-2.

43. Ramaswami, G. & Li, J. B. RADAR: A rigorously annotated database of A-to-I RNA editing. Nucleic Acids Research 42, D109–13 (2014).

44. Kiran, A. & Baranov, P. V. DARNED: A DAtabase of RNa editing in humans. Bioinformatics 26, 1772–1776 (2010).

45. Briese, M. et al. A systems view of spliceosomal assembly and branchpoints with iCLIP. Nature Structural and Molecular Biology 26, 930–940 (2019).

46. Picardi, E. & Pesole, G. REDItools: High-throughput RNA editing detection made easy. Bioinformatics (2013) doi:10.1093/bioinformatics/btt287.

47. Song, Y. et al. irCLASH reveals RNA substrates recognized by human ADARs. Nature Structural and Molecular Biology 27, 351–362 (2020).

48. Crews, L. A. et al. An RNA editing fingerprint of cancer stem cell reprogramming. Journal of Translational Medicine 13, 52 (2015).

49. Widmark, A. et al. ADAR1- and ADAR2-mediated regulation of maturation and targeting of miR-376b to modulate GABA neurotransmitter catabolism. Journal of Biological Chemistry 298, (2022).

50. Shannon, P. et al. Cytoscape: A software Environment for integrated models of biomolecular interaction networks. Genome Research 13, 2498–2504 (2003).

51. Bindea, G. et al. ClueGO: A Cytoscape plug-in to decipher functionally grouped gene ontology and pathway annotation networks. Bioinformatics 25, 1091–1093 (2009).

52. Ge, S. X., Jung, D., Jung, D. & Yao, R. ShinyGO: A graphical gene-set enrichment tool for animals and plants. Bioinformatics 36, 2628–2629 (2020).

53. Szklarczyk, D. et al. The STRING database in 2021: Customizable protein-protein networks, and functional characterization of user-uploaded gene/measurement sets. Nucleic Acids Research 49, D605–D612 (2021).

54. Quinones-Valdez, G. et al. Regulation of RNA editing by RNA-binding proteins in human cells. Communications Biology 2, (2019).

55. Castello, A. et al. Comprehensive Identification of RNA-Binding Domains in Human Cells. Molecular Cell 63, 696–710 (2016).

56. Beckmann, B. M. et al. The RNA-binding proteomes from yeast to man harbour conserved enigmRBPs. Nature Communications (2015) doi:10.1038/ncomms10127.

57. Zhan, X., Yan, C., Zhang, X., Lei, J. & Shi, Y. Structures of the human pre-catalytic spliceosome and its precursor spliceosome. Cell Research 28, 1129– 1140 (2018).

58. Cvitkovic, I. & Jurica, M. S. Spliceosome database: A tool for tracking components of the spliceosome. Nucleic Acids Research 41, (2013).

59. Murachelli, A. G., Ebert, J., Basquin, C., Le Hir, H. & Conti, E. The structure of the ASAP core complex reveals the existence of a Pinin-containing PSAP complex. Nature Structural and Molecular Biology 19, 378–386 (2012).

60. Lin, Y. C., Lu, Y. H., Lee, Y. C., Hung, C. S. & Lin, J. C. Altered expressions and splicing profiles of Acin1 transcripts differentially modulate brown adipogenesis through an alternative splicing mechanism. Biochimica et Biophysica Acta - Gene Regulatory Mechanisms 1863, (2020).

61. Rodor, J., Pan, Q., Blencowe, B. J., Eyras, E. & Caceres, J. F. The RNA-binding profile of Acinus, a peripheral component of the exon junction complex, reveals its role in splicing regulation. Rna 22, 1411–1426 (2016).

62. Raitskin, O., Cho, D. S. C., Sperling, J., Nishikura, K. & Sperling, R. RNA editing activity is associated with splicing factors in lnRNP particles: The nuclear pre-mRNA processing machinery. Proceedings of the National Academy of Sciences of the United States of America (2001) doi:10.1073/pnas.111153798.

63. Tariq, A. et al. RNA-interacting proteins act as site-specific repressors of ADAR2-mediated RNA editing and fluctuate upon neuronal stimulation. Nucleic Acids Research (2013) doi:10.1093/nar/gks1353.

64. Shanmugam, R. et al. SRSF9 selectively represses ADAR2-mediated editing of brain-specific sites in primates. Nucleic Acids Research 46, 7379–7395 (2018).

65. Hendrickson, D., Kelley, D. R., Tenen, D., Bernstein, B. & Rinn, J. L. Widespread RNA binding by chromatin-associated proteins. Genome Biology 17, 28 (2016).

66. Freund, E. C., et al. Unbiased Identification of trans Regulators of ADAR and A-to-I RNA Editing. Cell Reports 31, (2020).

67. Orecchini, E. et al. ADAR1 restricts LINE-1 retrotransposition. Nucleic Acids Research 45, (2017).

68. Hong, H. Q. et al. Bidirectional regulation of adenosine-to-inosine (A-to-I) RNA editing by DEAH box helicase 9 (DHX9) in cancer. Nucleic Acids Research 46, 7953–7969 (2018).

69. Shen, H. et al. ADARs act as potent regulators of circular transcriptome in cancer. Nature Communications (2022) doi:10.1038/s41467-022-29138-2.

70. Bonnal, S. C., López-Oreja, I. & Valcárcel, J. Roles and mechanisms of alternative splicing in cancer — implications for care. Nature Reviews Clinical Oncology vol. 17 457–474 at 10.1038/s41571-020-0350-x (2020).

71. Sahara, S. et al. Acinus is a caspase-3-activated protein required for apoptotic chromatin condensation. Nature (1999) doi:10.1038/43678.

72. Laurencikiene, J., Källman, A. M., Fong, N., Bentley, D. L. & Öhman, M. RNA editing and alternative splicing: The importance of co-transcriptional coordination. EMBO Reports (2006) doi:10.1038/sj.embor.7400621.

