## Supplementary figures and images for "ADAR1 regulates alternative splicing through an RNA editing-independent mechanism"

### Supp Figure 1

Supp. Figure 1

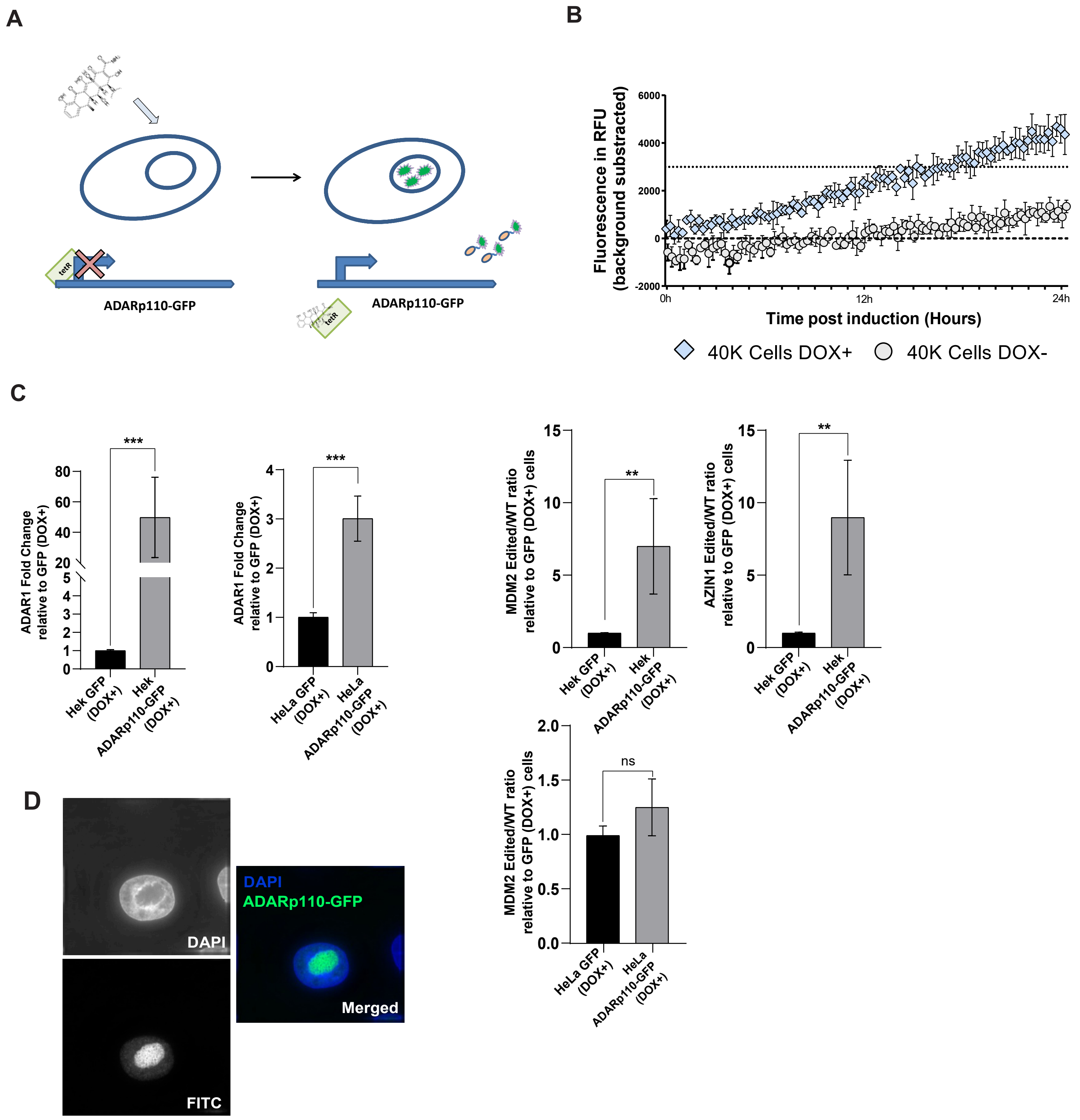

### Supp Figure 2

#### Supp. Figure 2

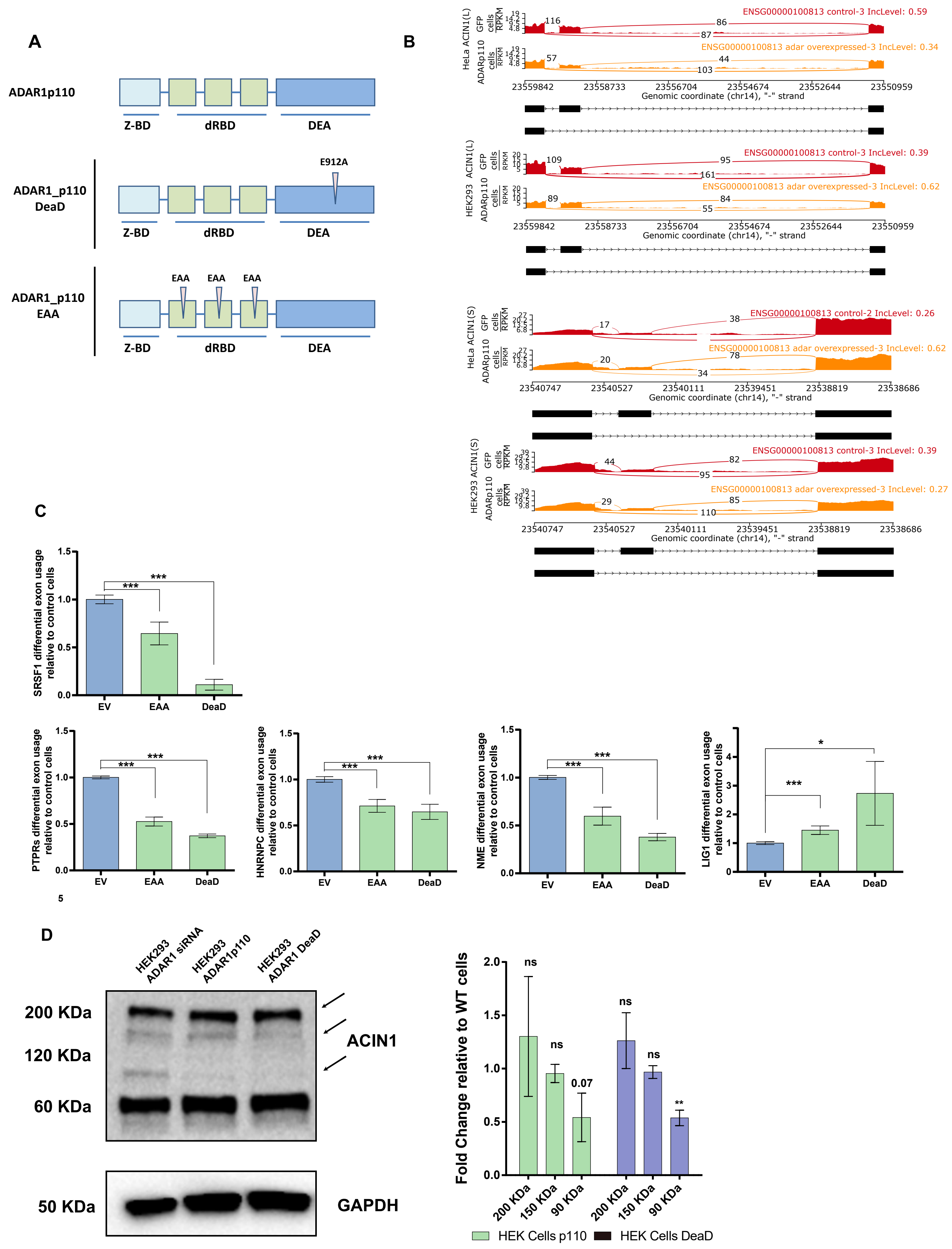

### Supp Figure 3

**Supp. Figure 3**

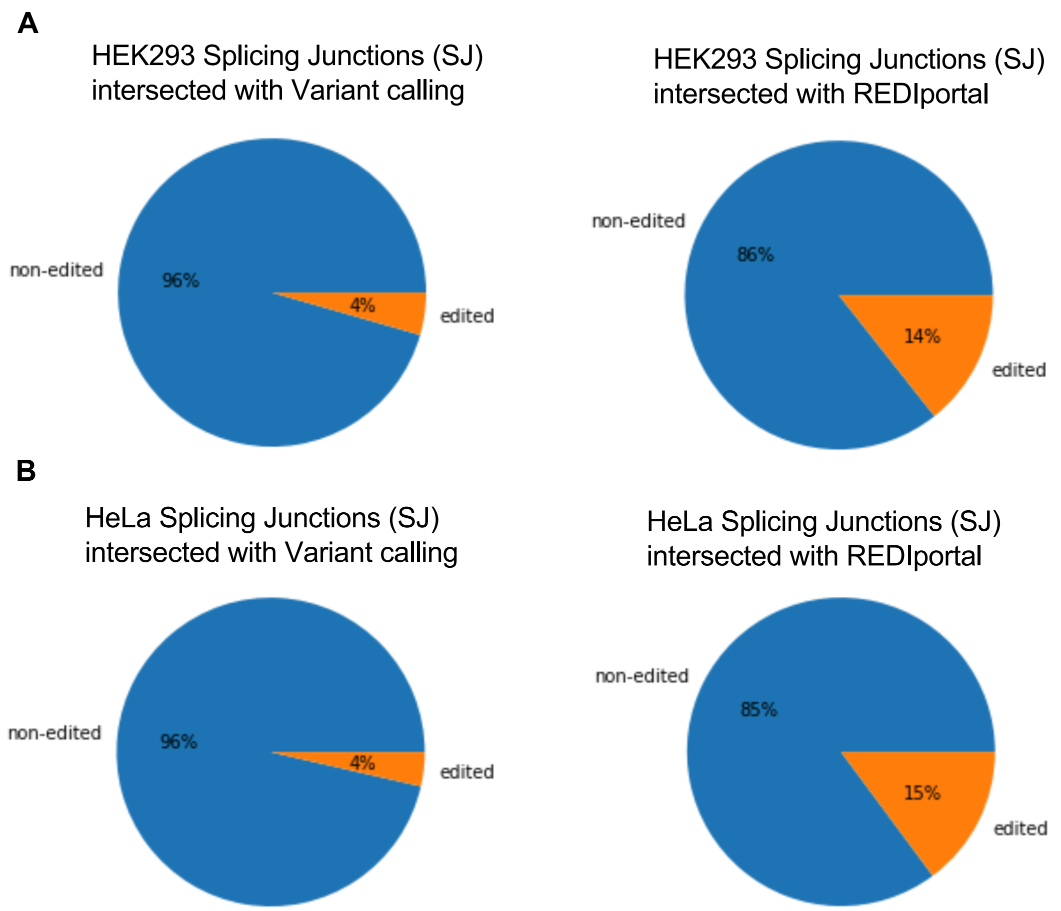
